# Elucidating Combinatorial Chromatin States by Co-ImmunoPrecipitation (comb-ChIP)

**DOI:** 10.1101/060962

**Authors:** Ronen Sadeh, Roee Launer-Wachs, Hava Wandel, Ayelet Rahat, Nir Friedman

**Affiliations:** School of Computer Science and Engineering and Institute of Life Sciences, The Hebrew University of Jerusalem, Jerusalem, Israel

## Abstract

Chromatin ImmunoPrecipitation followed by massively parallel sequencing (ChIP-Seq) has been instrumental to our current view of chromatin structure and function, and identifies correlating histone marks, which together demarcate biologically-relevant domains. However, as with most genome-wide assays, ChIP-seq is an ensemble measurement that reports on the average occupancy of individual modifications in a population of cells. Consequently, our understanding of the combinatorial nature of chromatin states relies almost exclusively on spatial correlations. Here, we report the development of a novel protocol, called indexed Combinatorial ChIP (comb-ChIP), which has the power to determine the genome-wide co-occurrence of histone marks at single nucleosome resolution. We show that at regions of overlapping ChIP signals, certain combinations of marks (H3K36me3 and H3K79me3) tend to co-occur on the same nucleosome, while other combinations (H3K4me3 and H3K36me3) do not, reflecting differences in the underlying chromatin pathways. We further use comb-ChIP to detect changes in histone mark co-occurrence upon genetic perturbation, illuminating new aspects of the Set2-RPD3S pathway. Overall, comb-ChIP promises to greatly improve our understanding of the structural and functional complexity of chromatin.

## Introduction

Nucleosomal histones, the fundamental packaging units of DNA, are massively decorated by a large number of posttranslational modifications (PTMs) or marks. These marks are highly conserved and play key roles in all genomic transactions (Rivera and Ren, 2013). Enzymes that deposit, remove, or bind histone marks are frequently mutated in human diseases such as cancer (Baylin and Jones, 2011; Chi et al., 2010; Maze et al., 2014). Chromatin ImmunoPrecipitation followed by next generation sequencing (ChIP-Seq) is used to determine the genome-wide location of nucleosomes bearing specific histone marks, and has been instrumental to our understanding of chromatin architecture, structure, and function in many cell types (ENCODE Project Consortium, 2012; Guttman et al., 2009; Weiner et al., 2015) ChIP-Seq studies in a variety of organisms identify combinations of spatially correlated histone marks; these combinatorial patterns demarcate biologically-relevant domains such as actively transcribed or polycomb repressed genes, heterochromatin, paused and active promoters, and enhancers, and can be used to predict unknown genomic functionalities (Guttman et al., 2009). However, a typical ChIP-Seq experiment reports on the average position and occupancy of a single modification at a time averaged over a large population of potentially heterogeneous cells. As a result, our current understanding of the combinatorial nature of chromatin states relies almost exclusively on spatial correlations between chromatin features. Biochemical studies have identified dozens of different histone marks, as well as multiple proteins that deposit, erase, and bind them. Surprisingly however, efforts to probe the complexity of chromatin in various cell types have identified a limited number of combinations of histone marks that specify defined genomic regions (ENCODE Project Consortium, 2012; Weiner et al., 2015). Mass spectrometry was proven to be a powerful tool for identification of histone marks complexity, however it lacks spatial information, and is limited to modification co-residing on a single, relatively short peptide (Garcia et al., 2007; Young et al., 2009). Recently, single molecule imaging allowed visualization of combinations of histone modifications, however it has limited spatial information (Shema et al., 2016). Importantly, it is generally unknown if spatially-correlated histone marks coexist or alternatively, if they represent different chromatin states occurring in different subsets of a population of cells (Figure 1A). Sequential ChIP, where histone marks are sequentially immunoprecipitated can report on the actual combinatorial nature of histone marks (Bernstein et al., 2006), yet such experiments are surprisingly rare and fraught with technical challenges. One factor which may impair the robustness and reliability of such experiments is large amount of input required to provide sufficient input to the second ChIP which can result in low signal to background ratio.

**Figure 1:**
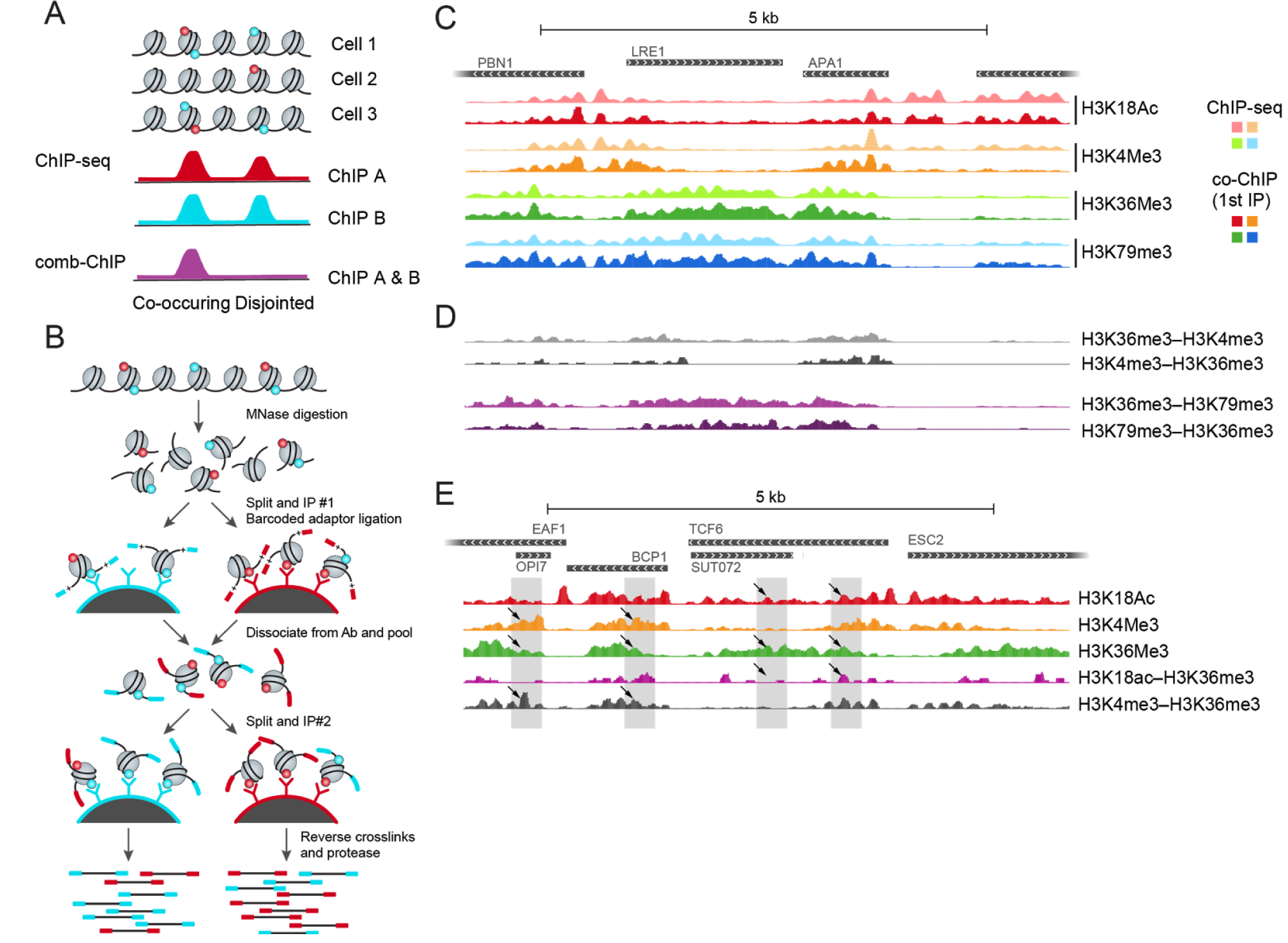
Comb-ChIP protocol for assaying combinations of histone modifications. **A** Overlapping signal of standard ChIP (red and cyan) can be due to co-occurrence of the two marks on the same nucleosomes (left peak, co-occurring), but can also be due to disjoint occurrence in the same location in different cells (right peak, disjointed). Combinatorial ChIP signal (purple) would allow to distinguish the two scenarios. **B** Outline of the comb-ChIP protocol: MNase digested chromatin is immobilized to magnetic beads coated with antibodies of interest (1st IP). Immobilized nucleosomes are ligated to barcoded adaptors to specify the 1st antibody. Following antibody inactivation and nucleosomes release, samples are pooled, redivided and subjected to a 2nd ChIP. Nucleosomes are reverse cross linked and NGS adapters are added by PCR to barcoded DNA to generate NGS-compatible libraries. At this stage a second barcode denoting the 2nd ChIP pool is added to the fragments. **C** The signal from the first ChIP step of the MNase comb-ChIP protocol (1st ChIP, solid colors) is in close agreement to standard MNase-ChIP (faded colors) (Weiner et al., 2015). Shown are coverage tracks for a representative genomic regions. **D** Reciprocal comb-ChIP signals are in good agreement. **E** comb-ChIP uncover co-occurrence and disjoint occurrences that are not available from individual ChIP: Shown is a representative genomic regions. The gray boxes highlight locations that are similar in terms of individual ChIP but different in comb-ChIP for these individual marks (black arrows).

Here, we report the development of a method, called Combinatorial ChIP (comb-ChIP) to map the genome-wide co-occurrence of histone marks at single nucleosome resolution (Figure 1B). Our starting point is the use of early barcoding (Lara-Astiaso et al., 2014; Rhee and Pugh, 2011; van Galen et al., 2016) and sample pooling (Lara-Astiaso et al., 2014). During the first immunoprecipitation step we ligate barcoded DNA adaptors to immobilized chromatin fragments. This barcoding enables pooling of ChIPed material prior to the second immunoprecipitation and preparation of NGS-compatible libraries. The barcoding and pooling solve two problems. First, by pooling we can multiplex many samples which allows us to use small amounts of input material per sample. Second, since the second IP is applied to multiple samples in a single tube, we reduce technical variability between samples. We provide experimental and analytical tools for efficient, reliable, and reproducible detection of combinations of histone marks.

### comb-ChIP can detect histone marks co-occurrence

To test the feasibility of chromatin barcoding for detecting coexistence of two histone marks, we selected well-established antibodies against promoter (H3K4me3, and H3K18ac) and gene-body (H3K36me3, and H3K79me3) histone marks. In comb-ChIP the first IP step is used for barcoding chromatin fragments, which provide the first layer of specificity (Figure 1B). Indeed, sequencing data obtained from our first IP and barcoding steps (input) are in good agreement with previously published traditional ChIP-seq datasets (Weiner et al., 2015) at both local (Figure 1C) and genomic (Figure S1) scales. To further determine whether this ChIP signal is specific, we repeated these assays in cells that express histones mutated at the antibody target residue (e.g., Histone 3 lysine 18 replaced by an arginine). The dramatic loss of ChIP material in relevant mutant background confirms that the barcoding step during the first ChIP is highly specific (Figure S2).

Following the first ChIP we pool the barcoded chromatin from the first ChIP and use it for the second IP step with the same battery of antibodies (Figure 1B), thus reading out pairwise existence of histone modifications. The use of MNase-digested chromatin ensures mononucleosome resolution for this assay. Comb-ChIP produced clear signal that was distinguishable from the parental ChIP experiments (Figures 1C,E), and was dependent on the integrity of the antibodies’ targets (Figure S2). Together, these observations suggest that the signal obtained by comb-ChIP is specific and is not the result of background interactions between the first and second ChIPs. Independent comb-ChIP experiments showed highly similar enrichment patterns (Figure S1B), Importantly, comb-ChIP signal was highly similar between reciprocal experiments in which the order of the two antibodies was reversed (Figures 1D and S1B) and exhibited different patterns than either of the relevant individual ChIPs (Figure S3), implying that comb-ChIP captures genomic location where the tested histone marks co-reside on a single nucleosome.

### comb-ChIP is a quantitative assay

We next turned to examine the quantitative nature of the comb-ChIP signal. At any genomic locus, we define the abundance of nucleosomes with a combination of two marks, (e.g., H3K36me3 and H3K79me3) as the fraction of cells in the population with a nucleosome with both marks at this locus. This abundance is naturally constrained by the abundance of individual marks at the same locus (Figure 2A). Specifically, the co-abundance of H3K36me3 and H3K79me3 cannot exceed the abundance of the individual marks, leading to two constraints (Figures 2B-C) on the co-abundance. On the other hand, if both the individual marks are highly abundant, than the co-abundance is constrained to be above a linear constraint (Figure 2D). These constraints hold for absolute abundances. However, the actual read counts in each comb-ChIP library depends on the abundance and also on other factors, such as antibody yield (fraction of targets retained in the IP step) and the sequencing depth.

**Figure 2:**
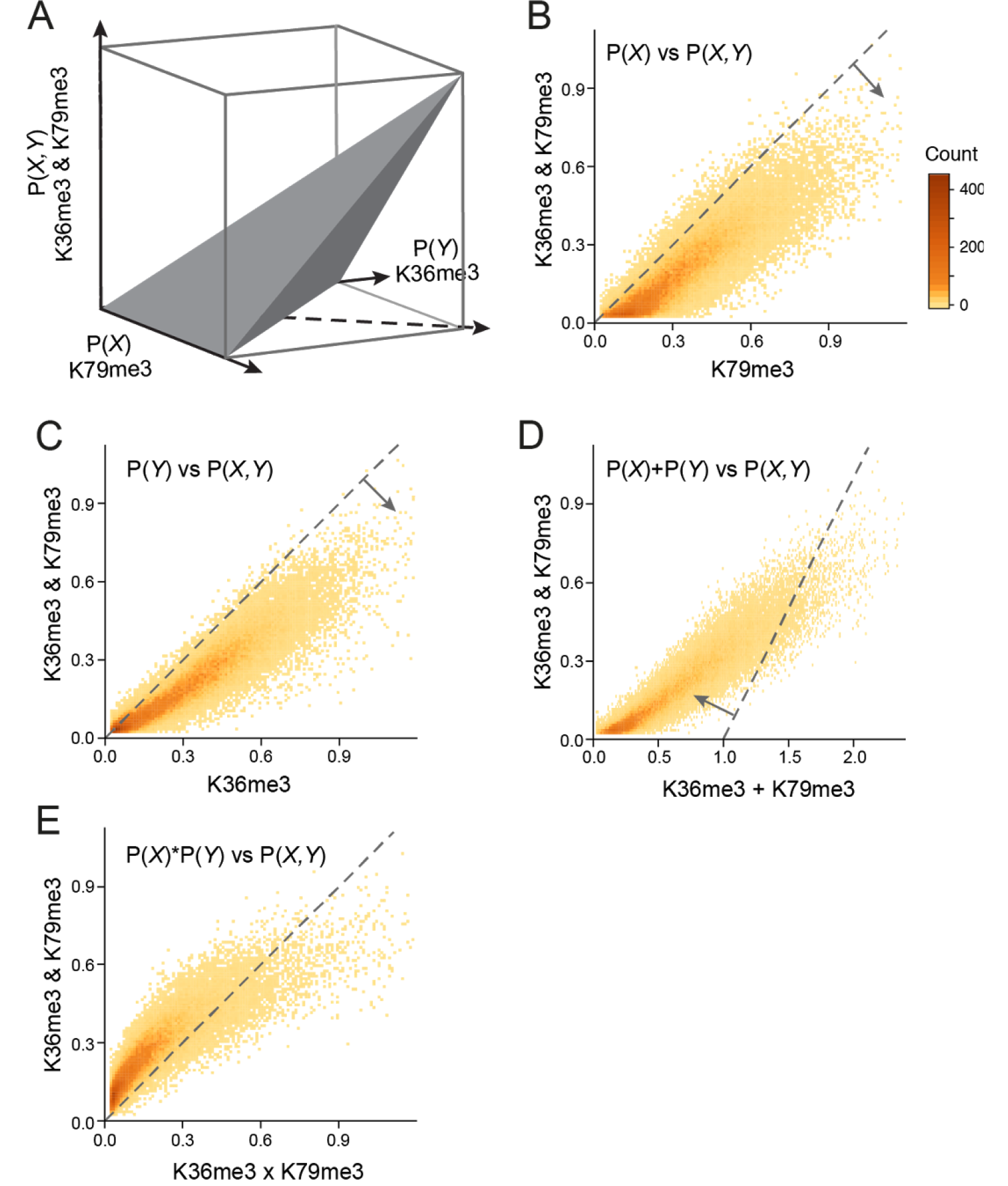
Comb-ChIP signal is a quantitative measure **A** Schematic description of the constraints relating the frequencies of individual marks at a location to the frequency of the dual marks at the same location. If comb-ChIP signal is quantitative then it should obey these constraints (up to a multiplicative constant). **B-D** Comparison of the individual ChIP of H3K36me3 and H3K79me3 to their comb-ChIP. Each panel interrogates one constraint (dashed lines). Occupancy levels are as assigned by the model’s transformation of the data (Methods). **E** Comparison of observed comb-ChIP signal against the value expected based on independence between the two individual marks.

We reasoned that if the signal is quantitative, we would expect to observe these constraints in the data up to an (unknown) amplification and measurement noise. In other words, the signal in each library should have a linear relation with the true abundances. This hypothesis leads to a testable prediction -- there is a multiplicative scaling coefficient that would make the comb-ChIP signal obey the underlying constraints. To test this prediction, we search for each comb-ChIP experiment for the scaling coefficient that minimizes the violations of these constraints (Figures 2B-D, Methods). Indeed, for each pair we find a scaling coefficient (one parameter) that agrees with the constraints (only up to 2.5% constraint violations) and spans the range of allowed interactions (Figures 2B-D and S4). Finding good scaling rules for our samples indicates that the comb-ChIP signal is approximately linear in the actual abundances. The scaled signal, for any given location, is an estimate of the abundance of combinatorial states in question at this locus, and thus provides a quantitative statement about each nucleosome location in each of the experiments. Specifically, this allows us to distinguish nucleosomes whose comb-ChIP signal is higher (or lower) than nucleosomes with similar predicted values (Figure 2E).

We observe that for most nucleosomes, the co-occurrence of H3K36me3 and H3K79me3 is higher than we would expect by independent model (Figure 2E). Both marks tend to accumulate in gene bodies in manner anticorrelated with nucleosome turnover rates(Dion et al., 2007; Venkatesh et al., 2012; Weiner et al., 2015). H3K36me3 is deposited by Set2, which is recruited by elongating RNA Pol II (Li et al., 2003), and its presence protects gene body nucleosome from eviction and thus reduces nucleosome turnover rates(Venkatesh et al., 2012). H3K79me3 is deposited by Dot1, in a manner that is dependent on H2B ubiquitylation (Ng et al., 2002). Moreover, there are no known histone demethylases that erase H3 lysine 79 methylation (unlike other lysine methylations). Thus, currently we assume that removal of H3K79me3 is only through nucleosome turnover. The comb-ChIP signal shows large co-occurrence of the two marks, even in nucleosomes with moderate levels of each individual mark. This observation is in agreement with the idea that H3K36me3 slows nucleosome turnover, which will result in accumulation of H3K79me3 on nucleosomes marked with H3K36me3.

### Co-occurrence of transcription associated marks

We next used this simple model to gain more insight into the relationship between H3K4me3 and H3K36me3. These marks are deposited by enzymes recruited by initiating (H3K4me3) and elongating (H3K36me3) forms of RNA Pol II that are differentially phosphorylated at the C-Terminal Domain (CTD) (Ng et al., 2003, 2002). Additionally, H3K36me3 has been implicated in suppressing transcription initiation from gene bodies (Carrozza et al., 2005; Venkatesh et al.,2012) . These suggest that H3K36me3 nucleosomes are prohibitive to transcriptional initiation and thus should be depleted of the initiation mark H3K4me3. This prediction is supported by the sparse overlap of the individual ChIP signals (Figure 3A). Examining the H3K4me4-H3K36me3 comb-ChIP values (Figure 3B,C), we see that about 50%-60% of the nucleosomes with noticeable signal for one of the marks do not display a comb-ChIP signal (12,019/23,064 and 10,025/30,221 of H3K4me3 and H3K36me3 nucleosomes, respectively). Focusing on the nucleosomes where both marks are present at the population level (Figure 3A, red outline), we observe comb-ChIP signal that is proportional to the expected values from a multiplicative model that assumes independence between the marks (Figure 3D). This is in contrast to the behavior of H3K36me3 and H3K79me3 (Figure 2E). Examining the location of nucleosomes with comb-ChIP signal for these marks we see that most of the comb-ChIP signal is in nucleosomes +3 to +5, which are in the overlap zone between the individual marks (Figure 3E). Moreover, this signal scales with expression level (Figure 3F). These results support a model where gene body nucleosomes have a low chance to be modified at both lysines during passage of any individual RNA Pol II molecule through the nucleosomes. Moreover, the border between the two modifications is fuzzy, either due to variation in timing of Pol II CTD modification or due to a large diameter of action of CTD-bound enzymes. Hence, the build up of both H3K4me3 and H3K36me3 on the same nucleosome is more likely to occur at highly expressed genes that experience repeated cycles of Pol II passages (Figure 3F).

**Figure 3:**
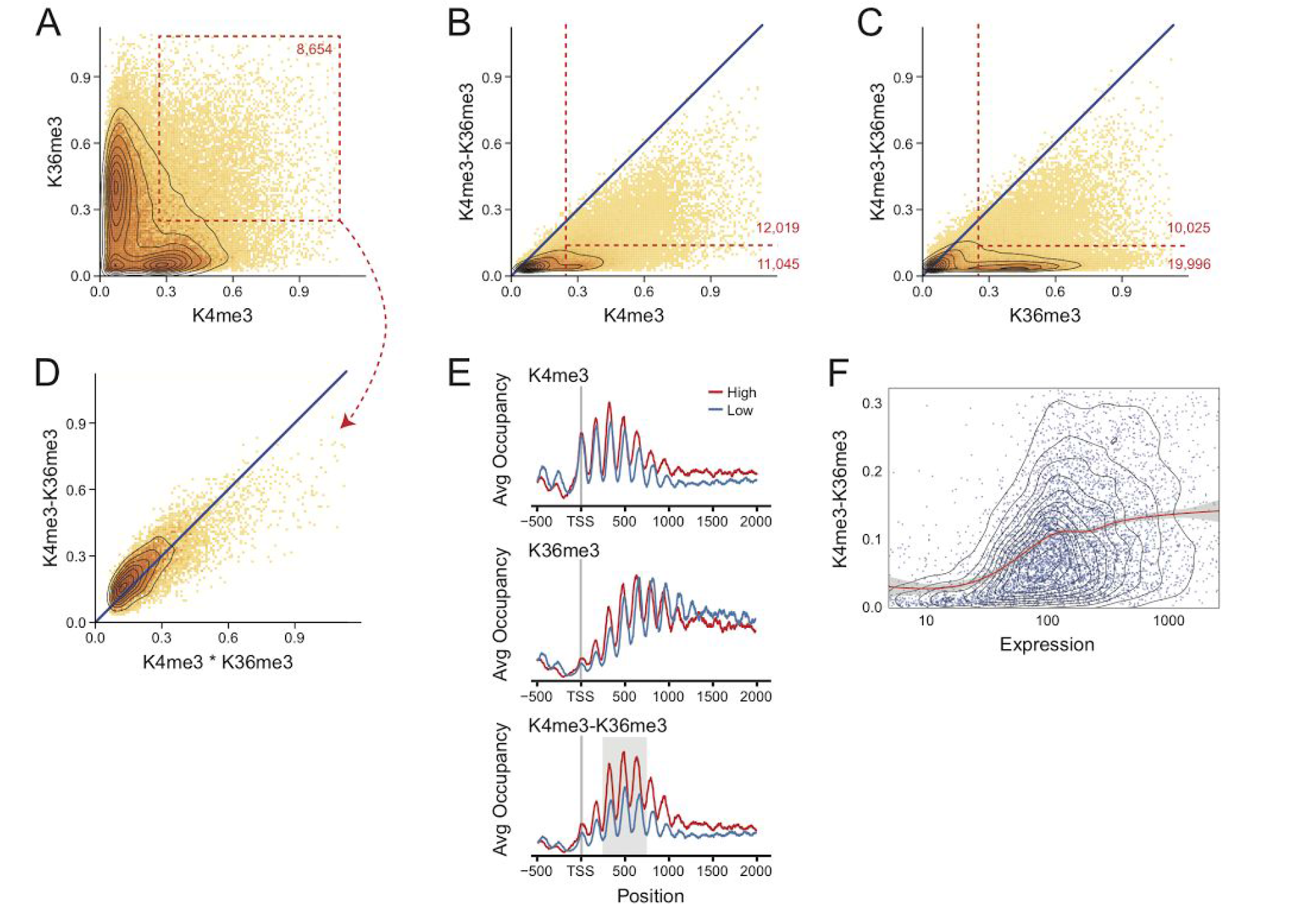
Co-occurrence of H3K4me3 and H3K36me3 scales with expression **A** Scatter of normalized H3K4me3 and H3K36me3 individual ChIP levels on all nucleosomes. Most nucleosomes have strong signal for one or the other marks. (red box) A subpopulation of nucleosomes with co-enrichment for both marks. **B-C** Comparison of individual ChIP to comb-ChIP (as in Figures 2B-C). Red lines denote population of nucleosomes with high levels of the individual ChIP signal and either low or high levels of comb-ChIP signal. Numbers in red denote number of nucleosomes in each region. B Comparison of expected combination by chance signal vs. comb-ChIP signal (as in Figure 2E) for the subpopulation marked in panel **B**. **E** Meta genes of ChIP signal of H3K4me3, H3K36me3 and their comb-ChIP. High, Low denote averages on genes in the 80%-100% and the 20-40% quantiles of expression, respectively. comb-ChIP signal is highest in nucleosomes +3-+5 (gray background). **F** Comparison of expression levels of genes to the average comb-ChIP signal on nucleosomes +3-+5 (area marked in gray in panel **E**). Red line marks the smoothed mean (gray area, confidence interval in the mean).

### Dissecting the Set2-RPD3S pathway

One of the best studied cases of chromatin regulation and crosstalk in yeast is the repression of cryptic transcription by the histone deacetylase Rpd3 small (RPD3S) complex (Figure S5). RPD3S is recruited to active gene bodies by RNA Pol II and likely gets activated by binding of its Eaf3 subunit to H3K36me3 (Drouin et al., 2010; Govind et al., 2010), which eventually lead to hypo-acetylation at gene body nucleosomes of active genes. As a result, interference with H3K36 methylation or RPD3S activity results in hyperacetylation of these nucleosomes, which in turn increases transcription initiation from “cryptic” promoters found in gene bodies (Carrozza et al., 2005; Venkatesh et al., 2012). These previous findings predict that Eaf3 knockout strains will have increased co-occurrence of H3K36me3 and H3K18ac at gene body nucleosomes. Indeed our assay reproduce previous reports (Carrozza et al., 2005; Venkatesh et al., 2012) showing increase in gene body H3K18 acetylation in cells lacking Eaf3 or Set2, with no apparent change in H3K36me3 in Eaf3 knockout cells (Figures 4A,B). We next tested the The comb-ChIP of H3K18ac and H3K36me3. As predicted, these marks comb-ChIP signal increases specifically at gene bodies in Eaf3 knockout cells (Figure 4C), thus directly probing this co-occurrence for the first time.

**Figure 4:**
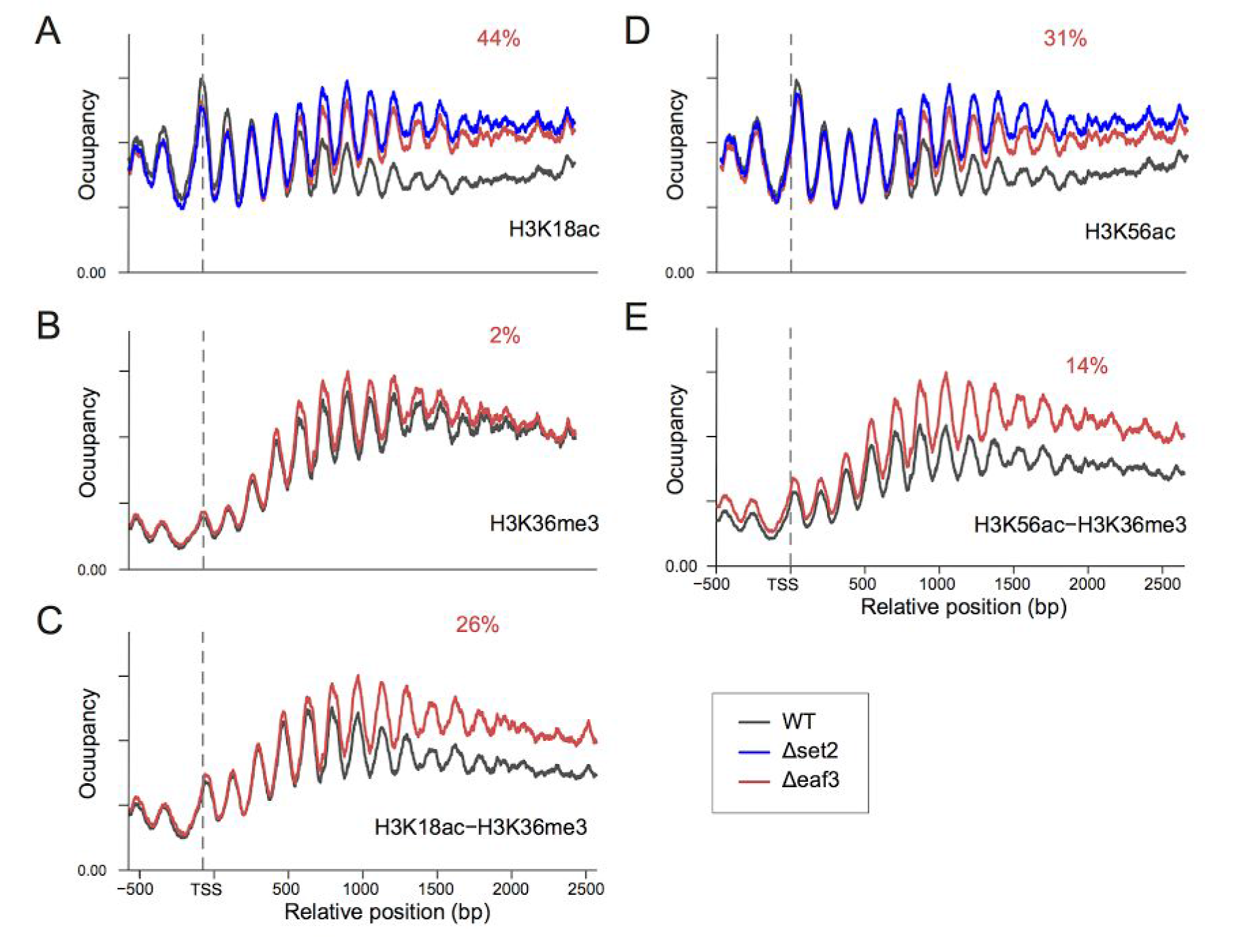
Gain of cooccurrence of marks as a result of genetic perturbation. Metagene profiles over long genes (ORF of 2000bp or longer). Average occupancy (arbitrary units) vs location relative to TSS in different strains. Median increase in gene body signal in Eaf3 knockout (relative to the increase in 5’ signal for each gene) is reported in red.

An additional mechanism by which H3K36me3 potentially reduces the acetylation level at gene-body nucleosomes is inhibition of nucleosome eviction by Pol II. Thus, in the absence of H3K36me3 mark, transcription evicts more nucleosomes. These are subsequently reassembled from newly synthesized histones (Venkatesh et al., 2012). Nascent histones are acetylated at several sites including H3K56 (Kuo et al., 1996; Masumoto et al., 2005), whose level is a proxy to nucleosome turnover rate(Rufiange et al., 2007) (Figure S5). Indeed, deletion of Set2 results in higher level of H3K56ac at gene-body nucleosomes (Figure 4D). Interestingly, we detect similar increase in H3K56ac in Eaf3 knockout cells (Figure 4D). This was quite surprising to us as the control of nucleosome turnover by H3K36me3 was linked to the Isw1 component Ioc4(Smolle et al., 2012). another H3K36me3 binding protein and not to Eaf3/RPD3S. It is possible that gene-body nucleosome hyperacetylation, observed upon Eaf3 deletion, reduces the stability of these nucleosomes, which increase turnover rate at these locations. We next tested the co-occurrence of H3K56ac with H3K36me3 (Figure 4E). We detect clear increase of the comb-ChIP signal for these marks at gene body nucleosomes similar to H3K18ac (Figure 4C). Since nascent histones are not known to be H3K36 tri methylated there are two possibilities that can lead to its co-occurrence with H3K56ac.1) Methylation of H3K36 takes place on recently assembled nucleosomes prior to their deacetylation by the H3K56ac-specific deacetylases Hst3/4.2) Unexpectedly, RPD3 might be able to deacetylate H356 upon exposure of this lysine due to partial disassembly of the nucleosome by Pol II. While it will highly interesting to tackle the mechanistic details of this unexpected co-occurrence, it demonstrates the ability of comb-ChIP to extend our current view of chromatin structure and function.

## Discussion

The experimental and computational framework presented in this manuscript present an important progress towards determining the genome-wide co-occurrence of histone modification at a single nucleosome resolution. We demonstrate the power of comb-ChIP in resolving population correlations of histone post translational modifications into functional understanding of chromatin states. Using comb-ChIP we were able to demonstrate that a certain combination histone marks (H3K36me3 and H3K79me3) tend to co occur while another (H3K4me3 and H3K36me3) shows random overlaps. These differences likely reflect underlying biological mechanisms that drive the co-accumulation of these histone marks. We further used comb-ChIP can shed new light on long lasting problems in chromatin biology. By applying comb-ChIP to cells compromised on the Set2-Rpd3 pathway we surprisingly find that a mark for newly assembled nucleosomes (H3K56ac) coexists with H3K36me3, which serve to inhibit nucleosome turnover.

Comb-ChIP provide a powerful extension of the widely used ChIP-seq assays. It is not limited to histone marks and can be readily adapted for detecting the co-occurrence of transcription factors as well as other chromatin-associated molecules. As with ChIP-seq, comb-ChIP relies on antibodies with the known caveats of antibody specificity and sensitivity. By distinguishing co occurring from merely correlating histone marks, comb-ChIP can help to pinpoint marks that co-specify distinct chromatin states (Ruthenburg et al., 2011).

In summary, MNase comb-ChIP is a robust, straightforward methodology that can be easily adjusted to robotic frameworks to probe the combinatorial nature of chromatin. We believe that comb-ChIP can greatly improve our understanding of the structural and functional complexity of chromatin.

## Methods

### Yeast strains

Yeast strains were obtained from the yeast KO collection (BY4741 with KanMX cassette replacing the deleted gene) and the histone substitution and deletion library (Dai et al., 2008). As WT strains we used Bar1 knockout from the yeast KO collection and the H3 WT from the histone substitution and deletion library.

### Cell growth, fixation, and MNase digestion

Yeast cells were grown in YPD media at 30°C with constant shaking to OD 0.6-0.8. Cells were fixed with 1% formaldehyde for 15 minutes at room temperature with occasional shaking, quenched with 0.125 M glycine for 5 minutes at room temperature with occasional shaking, collected by centrifugation, (4000 g, 5 minutes), washed with cold ddH2O supplemented with EDTA-free protease inhibitors cocktail (Roche) and the pellet was resuspended in buffer Z (1 M sorbitol, 50 mM Tris 7.4, 10 mM β-mercaptoethanol) with zymolyase (Seikagaku) at 0.3 - 1 units per 1 ml of original cell volume. Cells were gently rotated at 30°C for 25 minutes until > 95% of cells were spheroplasted. Spheroplasts were pelleted (6500 g, 10 minutes) and resuspended in NP buffer (10 mM Tris pH 7.4, 1 M sorbitol, 50 mM NaCl, 5 mM MgCl2, 1 mM CaCl2, and 0.075% NP-40, freshly supplemented with 1 mM β-mercaptoethanol, 500 μM spermidine, and EDTA-free protease inhibitor cocktail) at final concentration of 200 OD/ml. Chromatin was digested with 12.5 units/ml MNase (Worthington) for 20 minutes at 37°C, and digestion was stopped by removing the tubes into ice and addition of 1 volume of ice cold MNase stop buffer ( 220 mM NaCl, 0.2% SDS, 0.2% DOX, 10 mM EDTA, 2%,Triton X-100, EDTA-free protease inhibitor cocktail). Tubes were kept on ice for 10 minutes, vortexed 3 × 10 seconds, centrifuged (16,000 g, 10 minutes, 4°C), and the supernatant containing the nucleosomes was removed to a fresh tube.

### MNase digest evaluation

2-5% of the MNased chromatin was removed, treated with 1 μg RNase A for 30 minutes at 37°C, the volume was adjusted to 50 μl, SDS was added to 0.5%, and chromatin was treated with 50 units proteinase K for 2 hours at 37°C, and cross linking was reversed for 12-16 hours at 65°C. DNA was isolated by addition 2X SPRI beads, its concentration was measured by Qubit, and nucleosomes were visualized by TapeStation (Agilent). In all cases MNase pattern showed less than 80% mono nucleosomes to avoid over digestion.

### Chromatin immobilization

MNased chromatin equivalent to 100 ng of DNA as estimated by MNase digest evaluation was used per ChIP. Chromatin volume was adjusted to 100 μl with ice cold RIPA buffer (10 mM Tris pH 8.0, 140 mM NaCl, 1 mM EDTA, 0.1% SDS, 0.1% sodium deoxycholate, 1% Triton X-100, EDTA-free protease inhibitor cocktail) and antibody (for specific details see antibodies section below), and the samples were rotated for 2 hours at 4°C. 15ul of protein G dynabeads (washed three times in RIPA buffer) were added and samples were rotated for an additional hour. Samples were magnetized and the beads were washed 6 × RIPA buffer, 3 × RIPA 500 (RIPA containing 500 mM NaCI), 3 × LiCI wash buffer (10 mM Tris pH 8.0, 0.25 M LiCl, 0.5% NP-40,0. 5% Sodium Deoxycholate, 1 mM EDTA, EDTA-free protease inhibitor cocktail), 3 X 10 mM Tris pH 7.5.

### Chromatin barcoding and release

**End repair**: Immobilized chromatin was suspended in 20 μl of 10 mM Tris pH 7.4, and 40 μl of end repair mix [50 mM Tris pH 7.5, 10 mM MgCl2, 10 mM DTT, 10 mM ATP, 10 mM each dATP, dCTP, dGTP, dTTP, 0.375 units T4 polynucleotide kinase (NEB), 0.01 units T4 polymerase (NEB)] was added. Samples were mixed well, and incubated for 22 minutes at 12°C followed by 22 minutes at 25°C. Chromatin was washed once in 150 μl 10 mM Tris pH 8.0 and resuspended in 40 μl of 10 mM Tris pH 8.0.

**A base addition**: 20 μl of A-Base mix [10 mM Tris pH 8, 10 mM MgCl2, 50 mM NaCl, 1 mM DTT, 0.58 mM dATP, 0.75 units Klenow fragment (NEB)] was added to the beads, mixed well, and samples were incubated at 37°C for 30 minutes. Chromatin was washed once in 150 μl 10 mM Tris pH 8 and resuspended in 18ul of 10 mM Tris pH 8

**Adapters ligation**: 5ul of indexed adapters (Blecher-Gonen et al., 2013) were added to each sample, beads were mixed well and 34 ul of ligation mix [29 μl of 2X quick ligase buffer (NEB), 5ul quick ligase (NEB)] was added, beads were mixed well and incubated at 25°C for 45 minutes.

**Chromatin release:** This step releases bound chromatin and inactivates the antibodies used in the first ChIP. From this point it is important to keep samples at temperature higher than 15°C to prevent precipitation. Beads were resuspended in 12.5 μl freshly made 0.1 M DTT and incubated at RT for 5 minutes. 12.5 ul of freshly prepared 2X Chromatin Release Buffer (500 mM NaCl, 2% Deoxycholate, 2% SDS, 2 mM EDTA, 2X EDTA-free protease inhibitor cocktail) were added and the beads were incubated at 37°C for 45 minutes. Samples were pooled into a 1.5 ml tube, magnetize, the supernatant was removed into a fresh 1.5 ml tube, centrifuged max speed, 5 minutes, 15°C, and the supernatant was removed again to a 15 ml tube. Pooled samples were diluted by addition of 9 volumes of dilution buffer (100 mM Nacl, 10 mM Tris pH 8, 1 mM EDTA, EDTA-free protease inhibitor cocktail). Diluted samples were loaded on Amicon filter (Millipore UFC905024) (we usually load ~2 ml of diluted sample per Amicon filter) containing 12 ml of Amicon buffer (0.1% SDS, 0.1% Sodium Deoxycholate, 10 mM Tris pH 8,1 mM EDTA, 140 mM NaCl), and centrifuged at 2000 g, 20°C until ~ 0.25ml of concentrated sample is left in the filter. Concentrated samples were pooled together, 1 volume of Amicon equilibration buffer (2% Triton X-100, 0.1% SDS, 0.1% Sodium Deoxycholate, 10 mM Tris pH 8, 1 mM EDTA, 140 mM NaCl + 2X EDTA-free protease inhibitor cocktail) was added, and samples were vortexed. At this point samples can be flash frozen and stored at -80°C.

### Second ChIP and next generation sequencing

A critical point in this protocol is to use sufficient amount of barcoded chromatin from the first ChIP in the second IP step in order to end up with enough barcoded DNA for efficient library amplification. The amount of barcoded chromatin that should be used is dependent on factors such as antibody yield, modification abundance, and adapter ligation efficiency and should be determined empirically for each experiment. However, we find that pooling ~ 5 samples from the first ChIP gives good results in most cases.

The pooled barcoded chromatin was divided into fresh tubes according to the number of antibodies used for the second ChIP step. The volume was adjusted to 100 μl with RIPA buffer and the antibody, and chromatin immobilization and washes was done as for the first ChIP. Chromatin elution, and library amplification was done as described (Blecher-Gonen et al.,2013) . DNA libraries were paired end sequenced by Illumina NextSeq 500.

### Antibodies

The following antibodies were used in this study:

*For each antibody we used qPCR to determine the amount of antibody that results in the best yield /to background ratio.

| Antigen | Catlog # | µg antibody / ChIP |
| --- | --- | --- |
| H3 | ab1791 (Abcam) | 3 |
| H4 | 04-858 (Millipore) | 3 |
| H3K4me3 | 07-473 (Millipore) | 1 |
| H3K18ac | 07-354 (Millipore) | 3 |
| H3K36me3 | ab9050 (Abcam) | 3 |
| H3K79me3 | ab2621 (Abcam) | 1 |
| H3K56ac | 07-677 (Millipore) | 2 |

### Read Mapping

Pair-end reads were mapped to the yeast genome (sacCer3) using bowtie2 with maximal fragment size of 1000bp. We treated duplicate fragments as potential PCR artifacts. We thus treated the set of unique fragments found as the read-set. We defined mononucleosome fragments as these shorter than 220bp.

### Nucleosome coverage

We used the nucleosome location atlas defined by Weiner et al(Weiner et al., 2015). We measured nucleosome coverage by counting the number of fragments overlapping a window of size 50bp around the center of the nucleosome.

### Model Normalization

Consider two modifications, X and Y. Let *X_l_*, and *Y_l_* denote the event that a nucleosome at location l has either mark. Thus, the abundance in the population of each mark, or the combination is *P*(*X_l_*), *P*(*Y_l_*), and *P*(*X_l_, Y_l_*), respectively. From the laws of probabilities we have three constraints on these three entities:

I. *P*(*X_l_*) ≥ *P*(*X_l_, Y_l_*)
II. *P*(*Y_l_*) ≥ *P*(*X_l_, Y_l_*)
III. *P*(*X_l_, Y_l_*) ≥ *P*(*X_l_*) + *P*(*Y_l_*) − 1

These three constraints define the boundaries of the allowed region shown in Figure 2A. We assume that the number of reads, 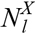,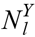 and 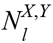 in our libraries are related to the abundance of each of the combination. The simplest assumption is that

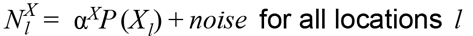

Similarly, the reads for other marks, each with its own multiplicative factor.

To test whether we can assign such multiplicative factor we did the following steps.

1. ChIP and comb-ChIP signals were divided by nucleosome occupancy, as measured by Weiner et al(Weiner et al., 2015)).
2. Each single ChIP nucleosome coverage vector was transformed to the range [0,1] by dividing by the 99.5% quantile value. This provides the single ChIP multiplicative factor.
3. For each comb-ChIP, we performed a line search for the scaling coefficient that minimizes the sum of the deviations from the allowed region constraints (as shown in Figures 2B-D). Formally, we define the loss of a factor α as

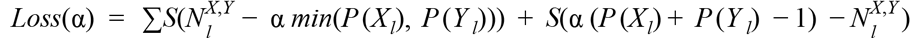

Where *S*(*z*) = *z* if *z* > 0, and 0 otherwise. The value of α that minimizes this loss is chosen for normalizing the counts for the comb-ChIP of *X* and *Y*

We repeated this procedure with different values of quantiles in Step 2. While the actual values were somewhat different, the relative conclusions, including differences from expected value (Figure 2E) were fairly robust to this choice.

### Density Plots and Smoothing

All plots were generated using the ggplot2 library of R (ver 3.2.3). Density scatters were generated using the geom_bin2d(), contours by geom_density2d() and smoothed averages by geom_smooth().

## Acknowledgements

We thank O.J. Rando and A. Appleboim for critical comments on the manuscript and L. Friedman and L. Gaffney for help with the graphics. This work was supported by ERC Grant 340712 and by the ISF I-Core on Chromatin and RNA in Gene Regulation.

---

**Supplementary Figure 1:**
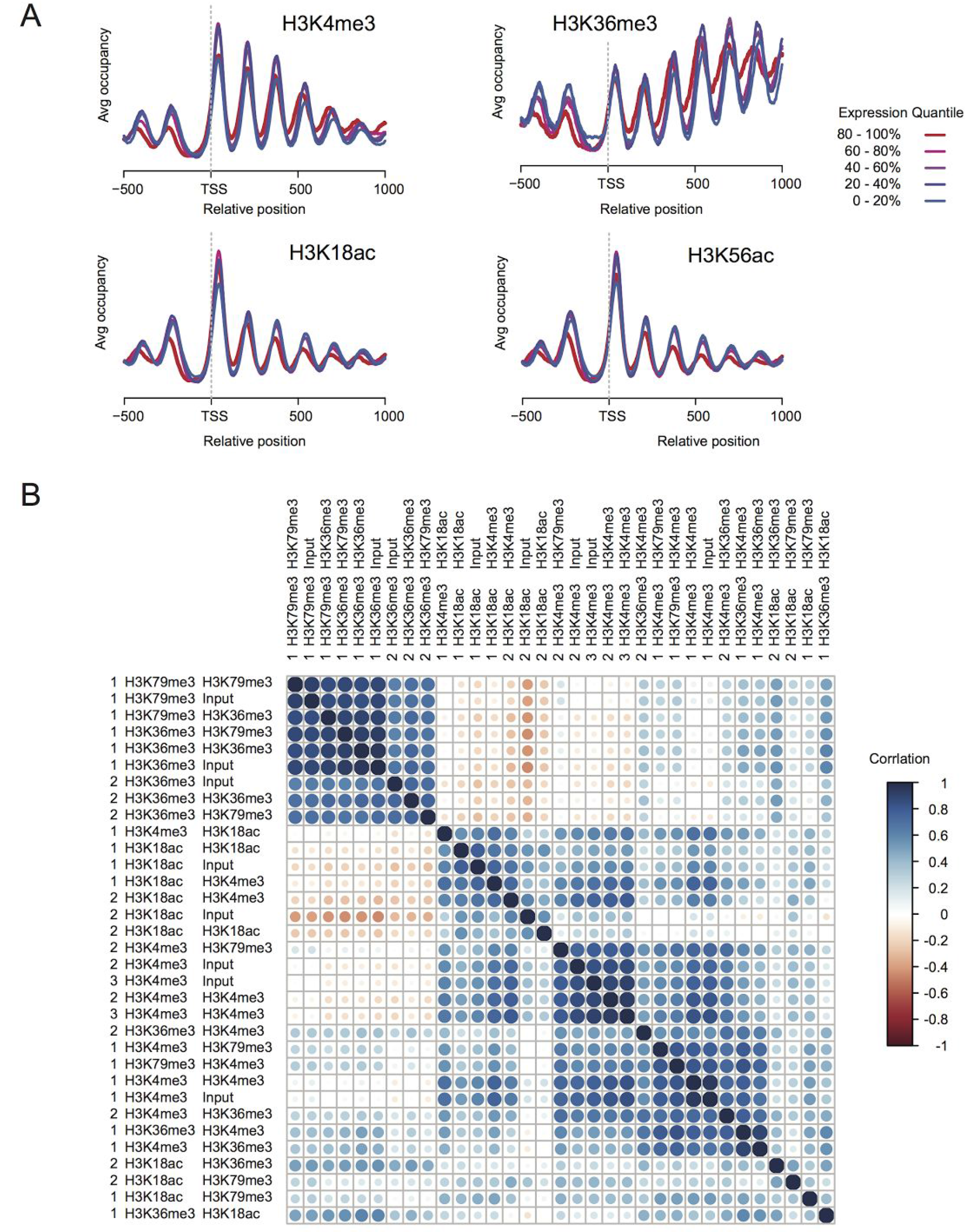
Comb-ChIP recapitulates standard ChIP-seq results and is reproducible. **A** Metagene profiles of the individual ChIP from our results (“input”). Each group of genes (sorted by expression quantiles) are averaged in TSS-aligned manner. **B** Correlation plot of nucleosome occupancy of different comb-ChIP tracks. Each name consists of batch number, first IP, and second IP (“input” first IP without second step). Correlation is denoted by color and by magnitude of the circle. Correlation was computed between coverage counts over nucleosomes (see Methods).

**Supplementary Figure 2:**
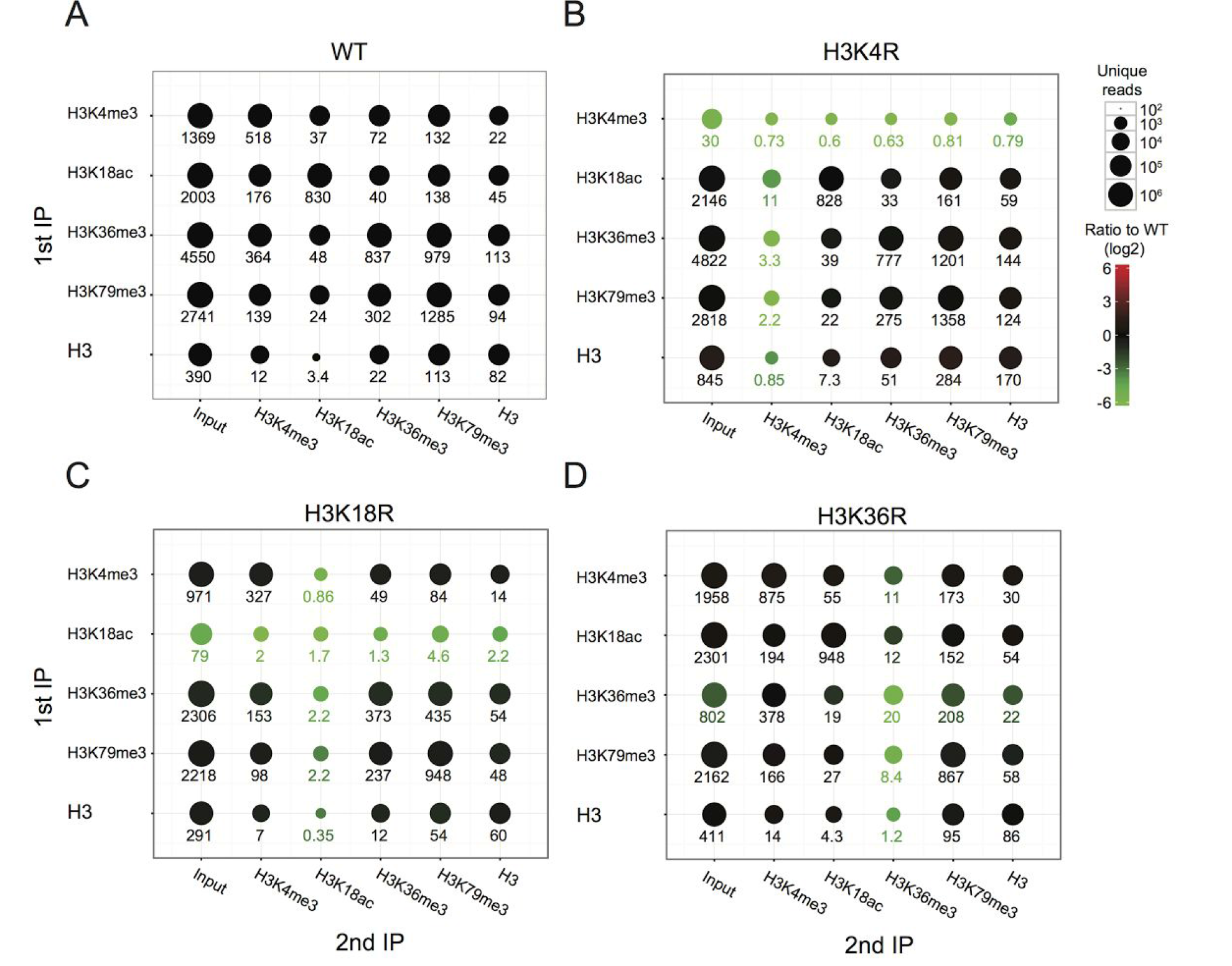
Comb-ChIP is specific. **A-D** Shown are the number of unique fragments recovered in experiment comparing pseudo-WT yeast to H3 mutants (Dai et al., 2008). Rows correspond to first IP and columns to second IP. The numbers of reads are shown in 1000s and denoted by the size of the circles. For each mutant, we denote the ratio to the corresponding comb-ChIP in WT by color scheme. For H3K4R and H3K18R we see significant reduction in the number of reads (~200-1000 fold). The effect is smaller in H3K36R due to cross reactivity of the specific batch of antibody used. Some of the imbalance in the reduction of read numbers between first IP and the second IP might be due to differences in tagging efficiency in different first IP steps.

**Supplementary Figure 3:**
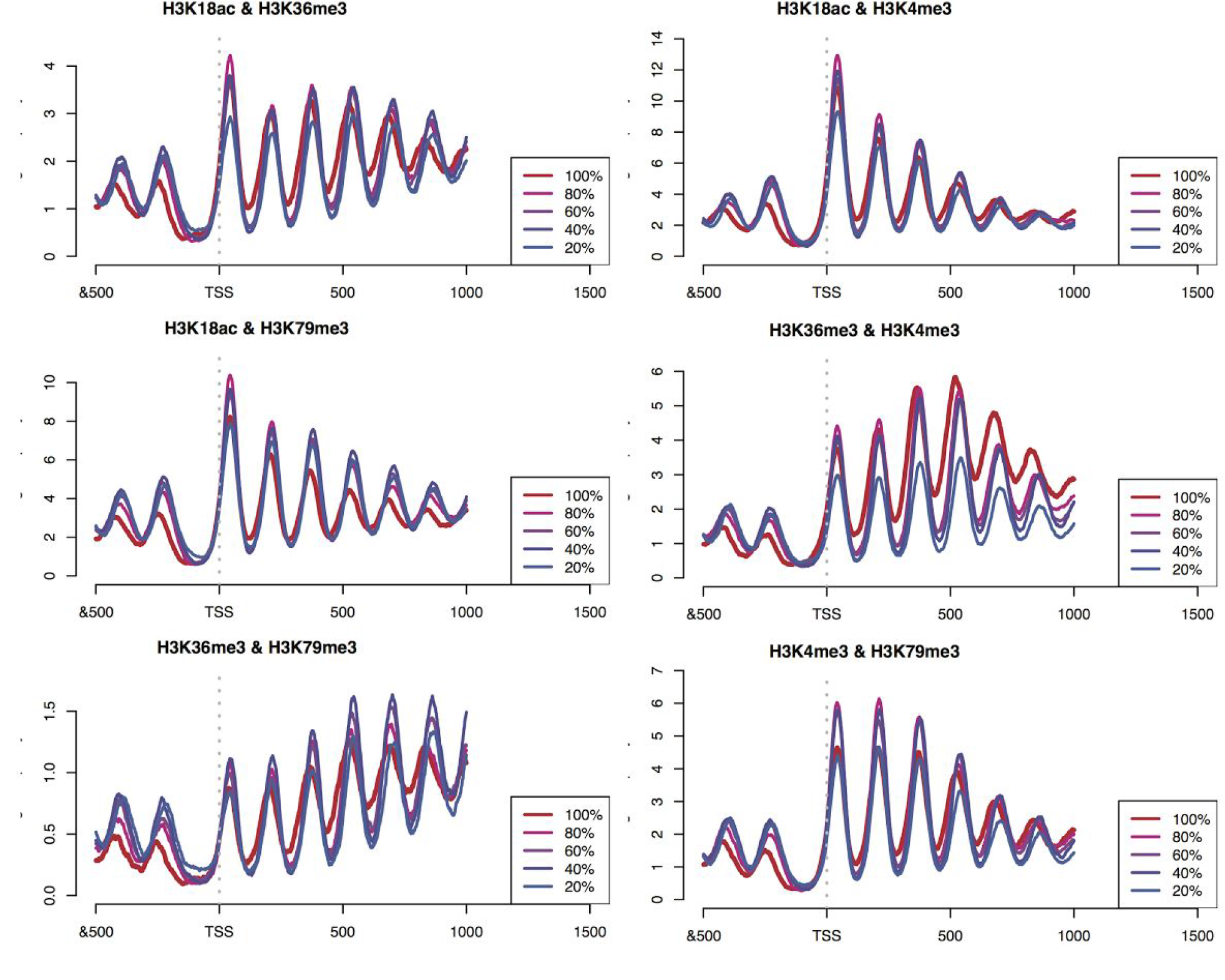
Comb-ChIP genic patterns Metagene profiles of comb-ChIP IP. Each group of genes (sorted by expression quantiles) are averaged in TSS-aligned manner.

**Supplementary Figure 4:**
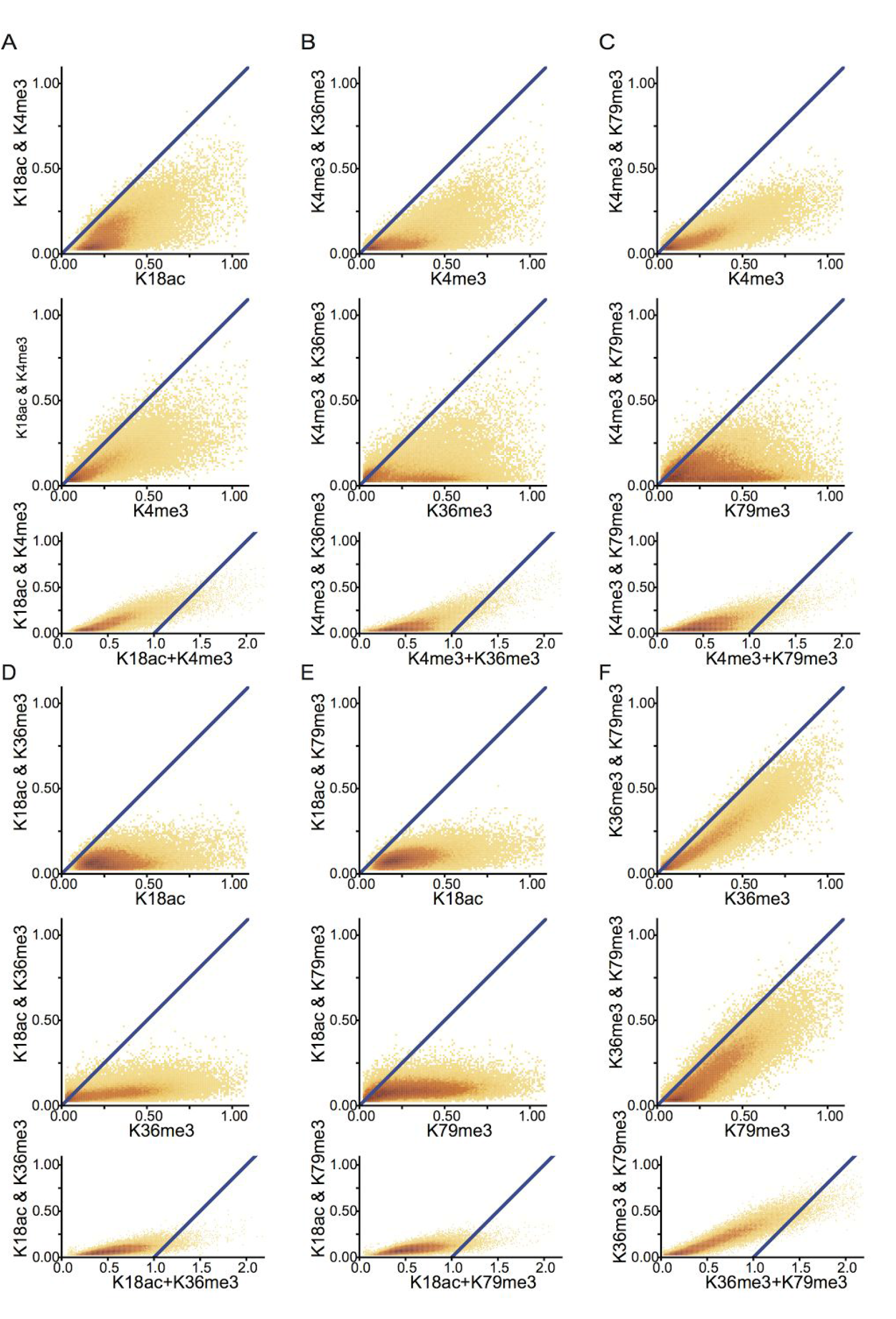
Quantitative comb ChIP patterns **A-F** Panels corresponding to Figure 2BD for each pair of marks.

**Supplementary Figure 5:**
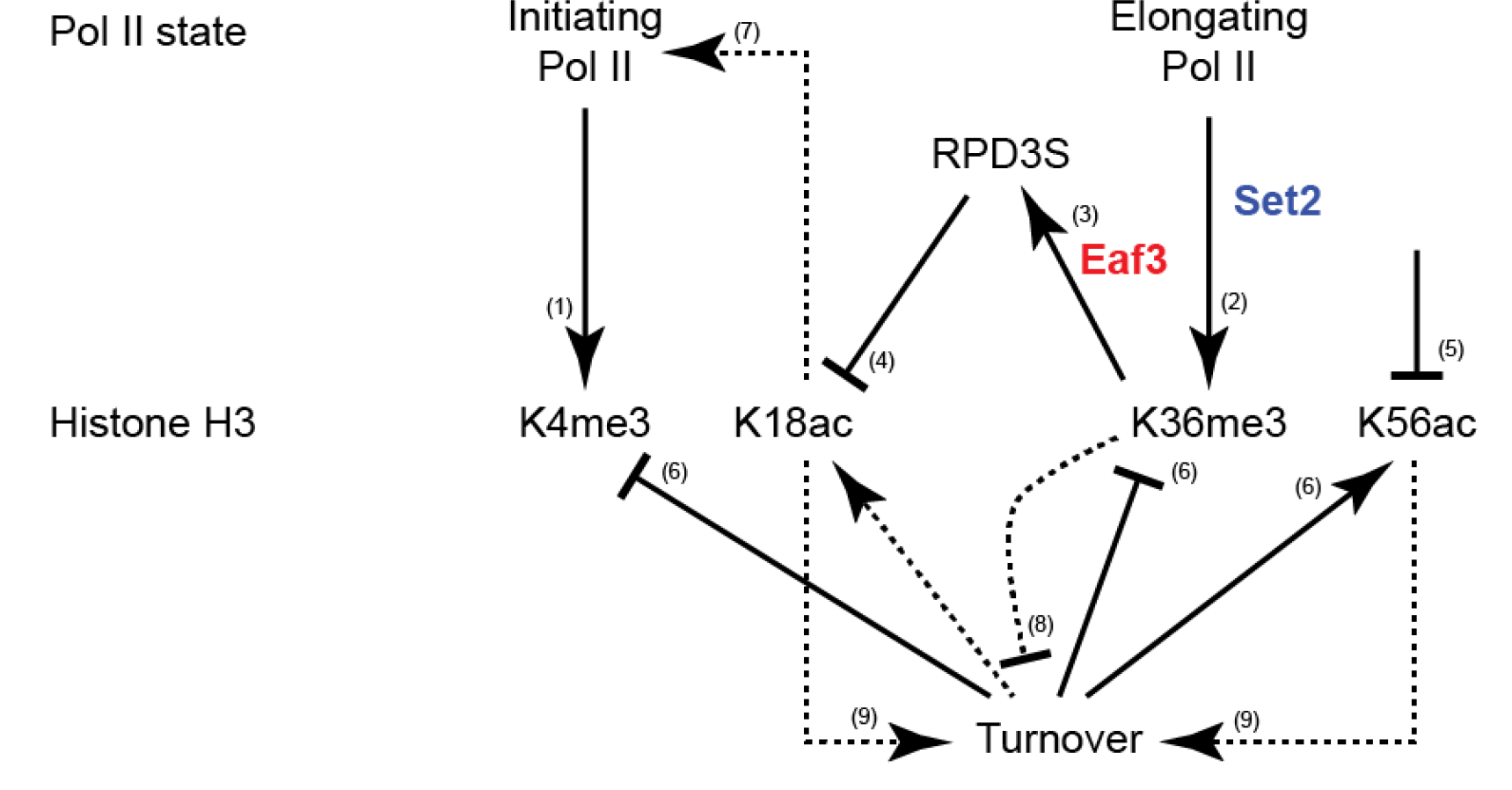
The Set2-RPD3 pathway Summary of the positive and negative relations between Pol II states, RPD3S recruitment, Histone H3 modifications, and turnover (Rando and Winston, 2012). Solid lines represent established connections: (1) H3K4me3 is deposited by Set1 recruited to initiating Pol II; (2) H3K36me3 is deposited by Set2 recruited to elongating Pol II (Li et al., 2003); (3) RPD3S is recruited to active gene bodies by RNA Pol II and likely gets activated by binding of its Eaf3 subunit to H3K36me3 (Drouin et al., 2010; Govind et al., 2010); (4) RPD3S deacetylates H3 tail lysines; (5) H3K56 is deacetylated by Hst3/4; (6) newly incorporated H3 is K56 acetylated and not tri-methylated in residues K4 and K36. Dashed lines represent indirect potential relations: (7) Deacetylated H3 nucleosomes repress transcriptional initiation; (8) H3K36me3 represses nucleosome turnover (Venkatesh et al., 2012); (9) Histone acetylation and specifically K56ac are associated with higher turnover.

